# Action-based organization and function of cerebellar cortical microcircuits

**DOI:** 10.1101/2020.04.04.025387

**Authors:** Nadia L Cerminara, Martin Garwicz, Henry Darch, Conor Houghton, Dilwyn E Marple-Horvat, Richard Apps

## Abstract

The cerebellum is the largest sensorimotor structure in the brain, but its mode of operation is not well understood. However, a fundamental organizational feature of the cerebellar cortex is division into elongated zones, defined by their inputs from specific parts of the inferior olive and Purkinje cell output to cerebellar and vestibular nuclei. Little is known about how the pattern of neuronal activity in zones, and their functional microcircuit subdivisions, microzones, is related to behaviour in awake animals. Here, we studied the organization of microzones within the C3 zone and their activity during a skilled forelimb reaching task in cats. Neurons in different parts of the C3 zone, functionally determined by receptive field characteristics, differed in their patterns of activity during movement. Our results suggest that the cerebellar C3 zone is organized and operates within an action-based frame of reference, with different microcircuits within the zone controlling specific muscle synergies.

## Introduction

A fundamental question in neuroscience is how neural information processing relates to behaviour. To be meaningful, any analysis of neural information processing needs to be based on a combined examination of anatomical connectivity and physiological activity in well-defined neural networks. By extension, therefore, the mapping of such structure-function relations is central to understanding the relationship between brain and behaviour.

The cerebellum is an ideal structure to examine such relationships. It has a well-defined cytoarchitecture (for reviews see Voogd and Glickstein, 1998; Ramnani, 2006; Apps and Hawkes, 2009; Cerminara et al., 2015) that has been studied extensively across macro-, meso- and microscale levels of resolution (e.g. Eccles et al., 1967; Palkovits et al., 1972; Palay and Chan-Palay, 1974; Brochu et al., 1990; Pichitpornchai et al., 1994; Wu et al., 1999; Serapide et al., 2001; Sugihara and Shinoda, 2004; Ke et al., 2009; Person and Raman, 2011; Stoodley et al., 2012; Heiney et al., 2014; Najafi et al., 2014; Proville et al., 2014; Yang and Lisberger, 2014; Giovannucci et al., 2017; Lesage et al., 2017; Marek et al., 2018; Streng et al., 2018; Aoki et al., 2019; Ashida et al., 2019; King et al., 2019). Its function has long been known to coordinate movements, but it is also thought to regulate autonomic, emotional and cognitive behaviour (Parsons et al., 2001; Strick et al., 2009; Schmahmann, 2010; Baumann and Mattingley, 2012; Buckner, 2013). The enormous neural information processing capacity of the cerebellum and its wide range of functions is reflected by the fact that it contains over half of all neurons in the brain (Herculano-Houzel, 2009), and it is extensively interconnected to the rest of the CNS (e.g. Anand et al., 1959; Snider and Maiti, 1976; Asanuma et al., 1983; Keifer and Houk, 1994; Brodal and Bjaalie, 1997; Teune et al., 2000; Kelly and Strick, 2003; Voogd and Ruigrok, 2004a; Suzuki et al., 2012; Carta et al., 2019; Watson et al., 2019). However, given the generally uniform cytoarchitecture of the cerebellum (although see Cerminara et al., 2015), it is widely assumed that the regulation of most, if not all, the functions that it serves relies on a ‘universal cerebellar transform’ (Ito, 1984; Schmahmann, 2010).

On a meso-scale, a key feature of the cerebellum is its division into a series of ‘modules’. Structurally, each module is defined by its climbing fibre input originating from a specific subdivision of the inferior olivary nucleus (e.g. Voogd and Bigare, 1980; Trott and Armstrong, 1987; Trott and Apps, 1991; Buisseret-Delmas and Angaut, 1993; Garwicz et al., 1996; Atkins and Apps, 1997; Voogd and Ruigrok, 2004b; Pijpers et al., 2006; Sugihara, 2011; Cerminara et al., 2013; Voogd et al., 2013).

Climbing fibres with a common olivary origin target one or more rostro-caudally oriented zones of Purkinje cells within the cerebellar cortex. These cortical zones are about 1-2 mm in mediolateral width, but traverse cerebellar lobules in the rostro-caudal direction for more than 10 mm (Oscarsson, 1979). In turn, Purkinje cells located within each cortical zone have a highly convergent projection to specific territories within the cerebellar and vestibular nuclei (Dietrichs, 1983; De Zeeuw et al., 1994; Garwicz et al., 1996; Apps and Garwicz, 2000; Voogd and Ruigrok, 2004b; Schonewille et al., 2006; Sugihara et al., 2009; Cerminara et al., 2013). These nuclei provide cerebellar output to spinal, brainstem and thalamic nuclei to influence behaviour. The cerebellum is thus divided into a series of olivo-cortico-nuclear modules which are highly conserved across species (Apps and Garwicz, 2005; Witter and De Zeeuw, 2015).

It is generally agreed that these modules are an important organizational feature of the cerebellum, with each module either acting individually or in combination with others to control behaviour (Ito, 1984; Apps and Garwicz, 2005; Manzoni, 2007; Cerminara and Apps, 2011; Witter and De Zeeuw, 2015; D’Angelo, 2018). However, high-resolution electrophysiological mapping within the cortical (zonal) component of individual modules has also revealed smaller units termed ‘microzones’. These are small groups of Purkinje cells arranged ~100-300 μm in the mediolateral axis that have similar climbing fibre peripheral receptive fields (Oscarsson and Uddenberg, 1966; Oscarsson, 1973; Andersson and Oscarsson, 1978; Andersson and Eriksson, 1981). The topography of microzones has been studied most extensively in the vermal B and paravermal C3 zones (Andersson and Oscarsson, 1978; Ekerot et al., 1991b; Hesslow, 1994; Jorntell et al., 1996).

Microzones and their associated olivo-cortico-nuclear microcircuits (collectively termed microcomplexes) are considered the fundamental operational unit of the cerebellum (Oscarsson, 1973; Andersson and Oscarsson, 1978; Oscarsson, 1979; Ito, 1984; Apps and Garwicz, 2005; Giovannucci et al., 2017). In relation to movement control individual microcomplexes are thought to control different aspects of the motor functions handled by each zone, but exactly what aspects of that function remain to be determined. Previous studies have examined the level of synchrony between Purkinje cells located within narrow, rostrocaudally oriented regions of cerebellar cortex that may correspond to individual microzones (Sasaki et al., 1989; Sugihara et al., 1993; Lang et al., 1999; Wise et al., 2010; Blenkinsop and Lang, 2011; Tang et al., 2019). More recently, this has been extended to studies in awake, head-fixed mice in relation to locomotion, spinal reflexes and also reward (De Gruijl et al., 2014; Kostadinov et al., 2019; Tsutsumi et al., 2019). However, these studies are confined to calcium imaging of Purkinje cell complex spikes and no information is currently available on Purkinje cell simple spike activity and cortical interneurons in relation to microzonal organization and skilled movement.

To address this issue, we have performed single unit electrophysiological recording from cerebellar cortical neurons with well-characterized receptive fields in the forelimb part of the C3 zone during the performance of a visually-guided forelimb reaching task in cats. Our experiments demonstrate that at the microzonal level: (i) different types of cortical neuron share receptive field characteristics; (ii) individually, neurons vary in their pattern of activity during task performance; (iii) however, Purkinje cell assemblies belonging to different receptive field classes encode different aspects of the broader motor function controlled by the cerebellar zone. These findings are consistent with an action-based organization and function of the cerebellar cortex.

## Results

### General features of the cerebellar neuronal population

Extracellular recordings were obtained from a total of 208 single units located in cerebellar lobules IV and V (Fig. 1A) within a region of paravermal cortex identified electrophysiologically as the forelimb part of the C3 zone (see Fig. 1B). Only single units with a clearly defined peripheral receptive field, located on the ipsilateral forelimb or shoulder (see Methods) were included in the analysis. A total of 65 single units met this strict criterion which allowed direct comparison with previous characterisation of C3 microzones in the anaesthetised cat (Ekerot et al., 1991a; Garwicz et al., 1998). These 65 units included 46 Purkinje cells (17 units with complex spikes, 38 units with simple spikes, for 9 Purkinje cells both complex spike and simple spike receptive fields were characterised, see Fig. 1C), 13 mossy fibres and 6 putative cortical interneurons (PCIs, see Methods for physiological identification of different cell types). Most of these units were found to be activated either by brushing of hairs or light tapping of the skin of the ipsilateral forelimb. Some units required pressure and palpation of deeper tissue within the forelimb, although none were found to be activated by palpation of individual muscles.

**Figure 1.**
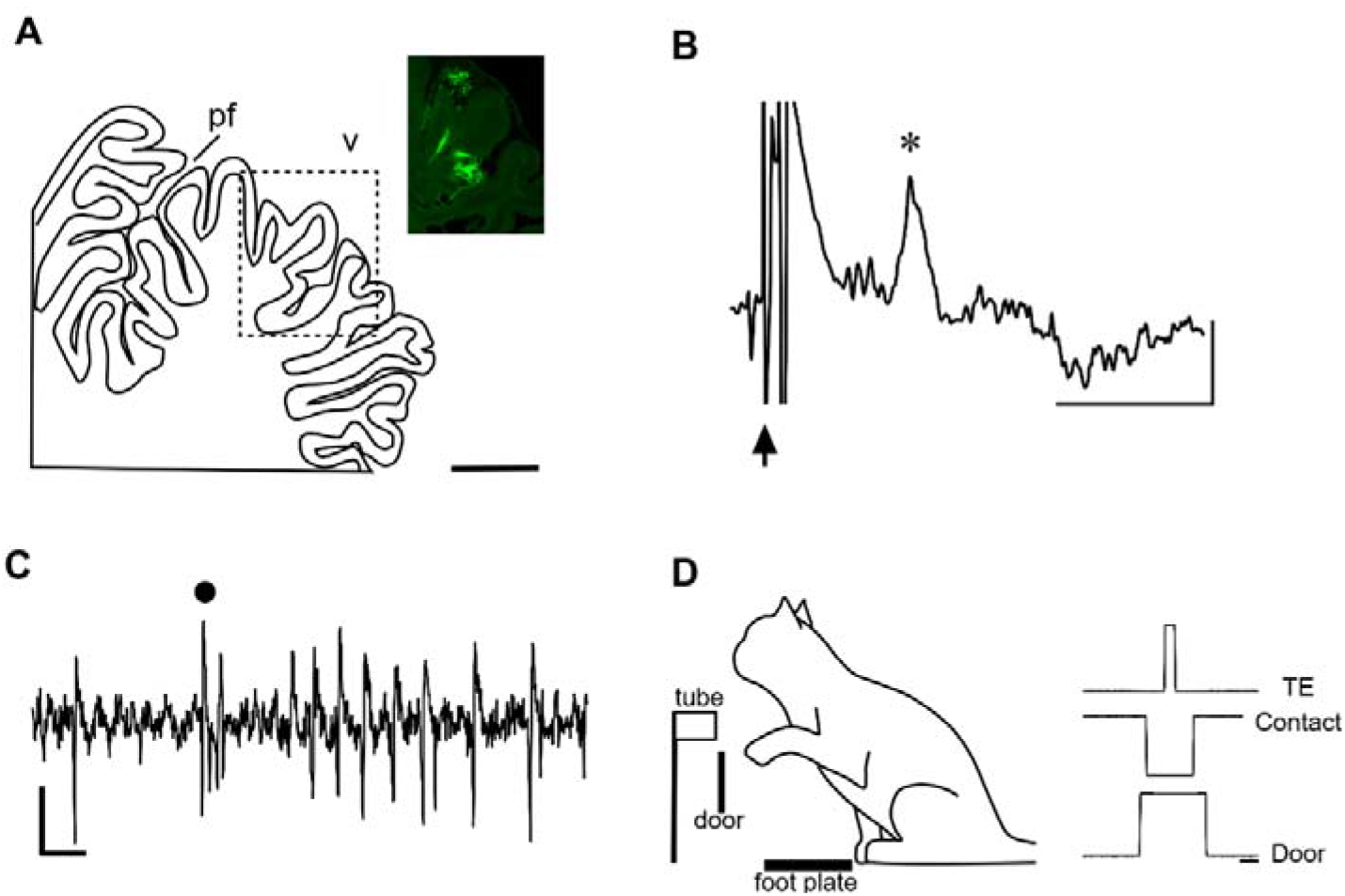
Experimental arrangements. **(A)** Sagittal outline of cerebellar section. Dotted rectangle shows area related to photomicrograph with fluorescent material in lobule V to mark the location where the majority of recordings where made. Pf, primary fissure; V, lobule V. Scale bar, 2 mm **(B)** C3 zone climbing fibre field potential (asterisk) evoked by ipsilateral superficial radial nerve stimulation (onset of stimulus at time of arrow, intensity, 2T). Scale bars, 50 μV, 10 ms **(C)** Example extracellular recording of a Purkinje cell in the C3 zone. Complex spike waveform marked by a dot. Scale bars, 50 μV and 5 ms **(D)** Schematic of reach retrieval task. The cue to reach was a door covering the tube dropping down. A contact foot plate monitored paw lift-off. At the end of the reach, the paw enters the tube, thereby breaking an infrared beam located at the mouth of the tube. Sequence of event markers recorded during a single reach retrieval are shown on the right. Door: upward deflection of trace represents tube door opening; Contact: downward deflection represents paw lift-off from contact plate whereas upwards deflection represents paw touch-down after retrieval of a food reward in the tube; TE: tube enter, upward deflection paw entering tube. Scale bar, 0.5 s.

### Topographical organization of C3 microzones in the awake animal

The topographical organization of Purkinje cell climbing fibre receptive fields within the C3 zone in cerebellar lobules IV and V, as determined previously in the anaesthetized cat is shown in Figure 2. As a first step, it was important to establish whether this arrangement was similar in the awake animal. In the awake cat, Purkinje cells with climbing fibre receptive fields on proximal and ventral areas of the ipsilateral forelimb were located in rostral folia of lobule V, while receptive fields restricted to the ipsilateral hindlimb were located further rostrally (figurines with receptive fields shaded dark blue in Fig. 2). The peripheral receptive fields of units identified as mossy fibres, PCIs or the simple spike activity of Purkinje cells could also be fitted to a similar spatial organization (figurines with receptive fields shaded light purple, pink or purple, respectively in Fig. 2). Therefore, regardless of neuronal type, the topographical organization of cerebellar cortical units located within the C3 zone in the awake cat was consistent with the detailed mapping previously reported in the anaesthetised animal.

**Figure 2.**
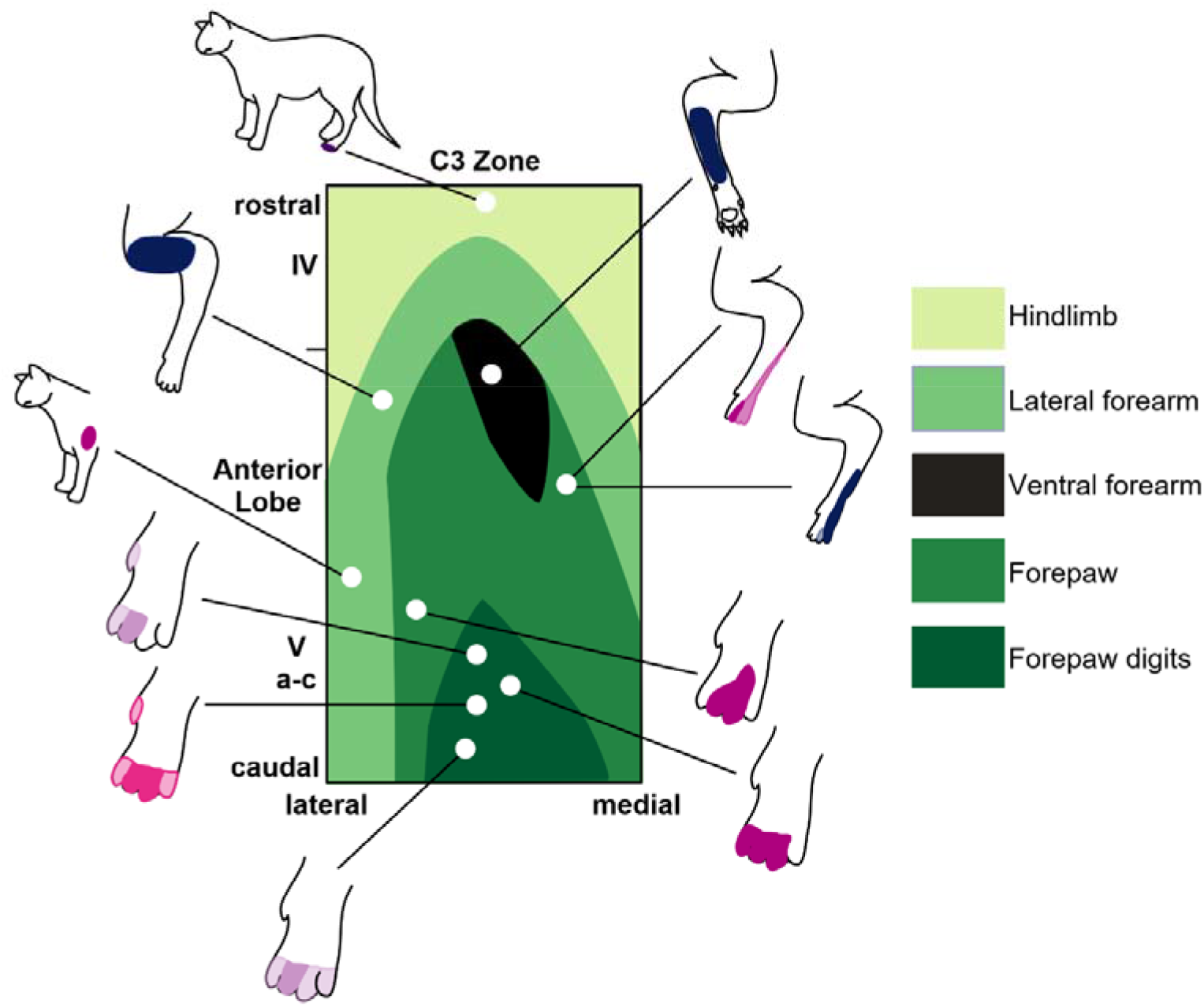
Topographic organization of the C3 zone in the awake cat. Map showing limb receptive field locations within the C3 zone in the anterior lobe as determined in the anaesthetised cat (Ekerot et al., 1991a). Cerebellar lobules are indicated to the left. Figurines surrounding the map show example peripheral receptive fields in the awake cat recorded in different recording sessions from two animals. Figurines with receptive fields in dark blue are Purkinje cell complex spikes, in purple are Purkinje cell simple spikes, in light purple mossy fibres and pink cortical interneurons. Darker shading in figurines denotes regions of skin that generated the strongest response within the receptive field; lighter areas denote total extent of receptive field.

### Forelimb receptive field classification

Receptive field data were available for all but one of the 10 receptive field classes (classes 1-6, 8-10) defined previously in the anaesthetised animal. Figure 3 (see also Supp. Fig. 1 for receptive fields of individual units) shows for each of these classes the total area of receptive field found for each neuronal type (Purkinje cell complex spikes, Purkinje cell simple spikes, mossy fibres and PCIs). Overall, the receptive fields covered most parts of the ipsilateral forelimb, apart from the ventral surface of the upper forelimb. Class 1 units were those with receptive fields located on the ventral and dorsal surface of the digits (Fig. 3A, n=13 units). Class 2 units were those with receptive fields (Fig. 3B, n=6 units) located on the ventral and dorsal surface of the forepaw, whereas class 3 units (Fig. 3C, n=6 units) had receptive fields that were located predominantly on the dorsal paw and forearm. Class 4 units (Fig. 3D, n=4 units) were those with receptive fields located predominantly on the ventral surface of the paw. Class 5 (Fig. 3E, n=2 units) were confined to the ventral surface of the paw and wrist or forearm while receptive fields located on the lateral part of the forearm were identified as class 6 (Fig. 3F, n=9 units). No cells were found with receptive field characteristics consistent with class 7 (i.e. lateral forearm and upper arm receptive fields). Class 8 cells were those with receptive fields located on the lateral shoulder/upper arm and neck (Fig. 3G, n=10), whereas units with receptive fields located on the radial/medial part of the paw and forearm were class 9 (Fig. 3H, n=17 units). Finally, class 10 cells (Fig. 3I, n=7 units) were those with receptive fields mainly confined to the medial shoulder/upper arm (Ekerot et al., 1991a; Jorntell et al., 1996; Garwicz et al., 1998).

**Figure 3.**
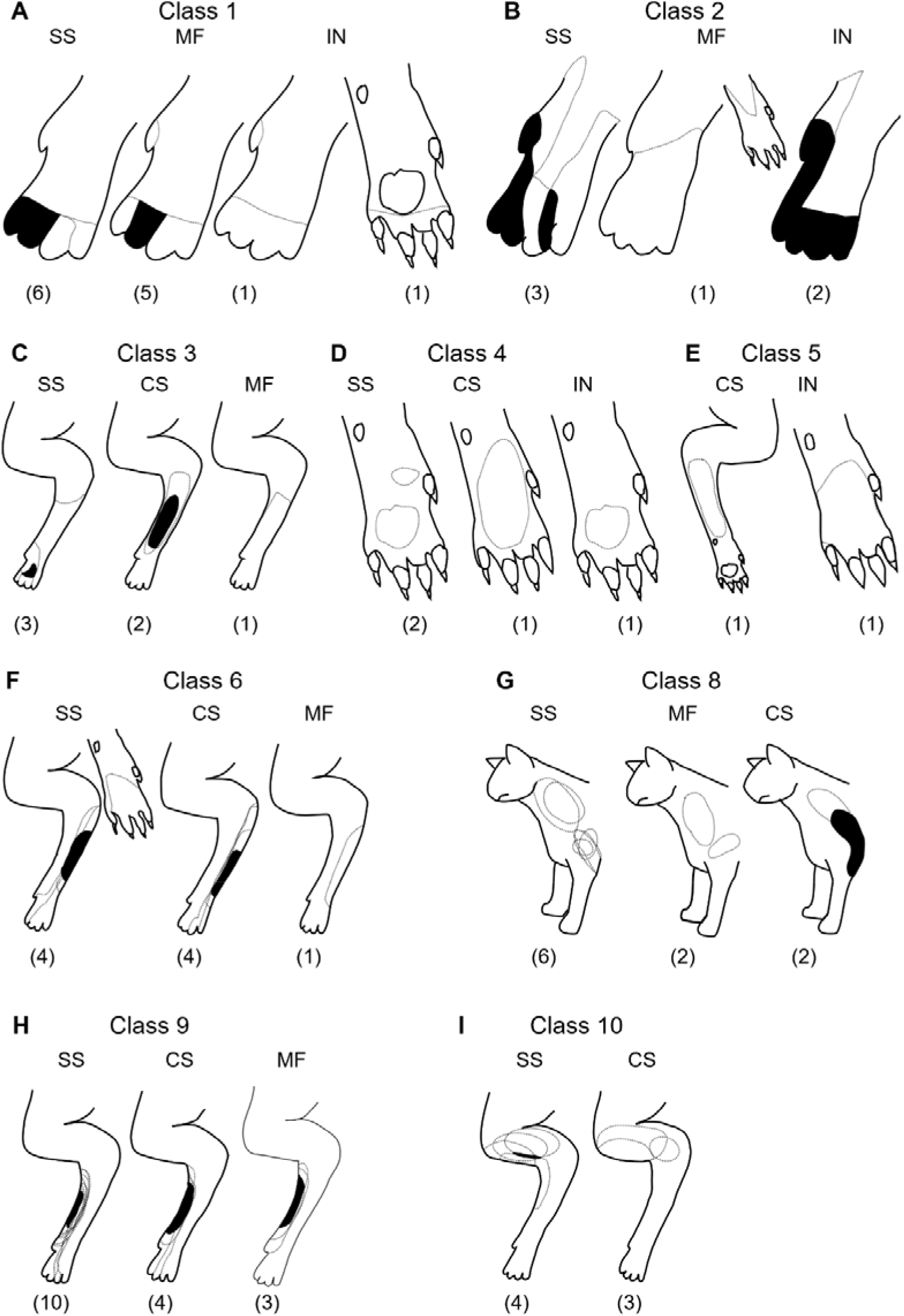
Classification of cerebellar cortical units according to receptive field class. Units were categorised in accordance with previous classification of Purkinje cell complex spike (Ekerot et al., 1991a) and mossy fibre receptive field characteristics (Garwicz et al., 1998). Each figurine shows all available receptive fields superimposed for each class and subdivided into those units identified as Purkinje cell simple spikes (SS), complex spikes (CS), mossy fibres (MF) or putative cortical interneurons (IN). Numbers in parentheses below each figurine indicate the number of units per type. **(A)** Class 1; **(B)** Class 2; **(C)** Class 3; **(D)** Class 4; **(E)** Class 5; **(F)** Class 6; **(G)** Class 8; (H) Class 9; (I) Class 10. Dashed lines indicate total extent of single receptive fields while areas filled in black indicate where overlap occurred in every available case.

### Comparison of receptive fields of simple spikes and complex spikes from the same Purkinje cells

An important disagreement in the literature is whether the simple spike receptive fields of Purkinje cells are similar to their complex spike receptive fields and therefore generated primarily due to local granule cells (‘local’, Bower and Woolston, 1983; Cohen and Yarom, 1998; Brown and Bower, 2001; Isope and Barbour, 2002), or differ and therefore generated mainly by non-local granule cells (‘non-local’, Ekerot and Jorntell, 2001; Jorntell and Ekerot, 2002; Ekerot and Jorntell, 2003; Dean et al., 2010). For nine Purkinje cells we were able to reliably discriminate simple spikes and complex spikes from the same Purkinje cells and define their corresponding receptive fields (Fig. 4). Receptive fields of complex spikes and simple spikes in six Purkinje cells (67%) were found to have similar peripheral receptive fields. Maximum overlap between complex spike and simple spike receptive fields was on average 64.4% ± 35.6 (n=6, pairs of receptive fields outlined in green in Fig. 4). The remaining three cells differed markedly in the location of their complex spike and simple spike receptive fields and had, on average, a maximum overlap of only 2.2% ±3.8 (n=3, pairs of receptive fields outlined in red in Fig. 4). Thus, the simple spike receptive fields of the majority of Purkinje cells in our sample were consistent with their activity being mainly (if not exclusively) the consequence of excitation of local granule cells. The exceptions suggest, however, that in some instances simple spike activity may be dominated by excitation from granule cells located non-locally.

**Figure 4.**
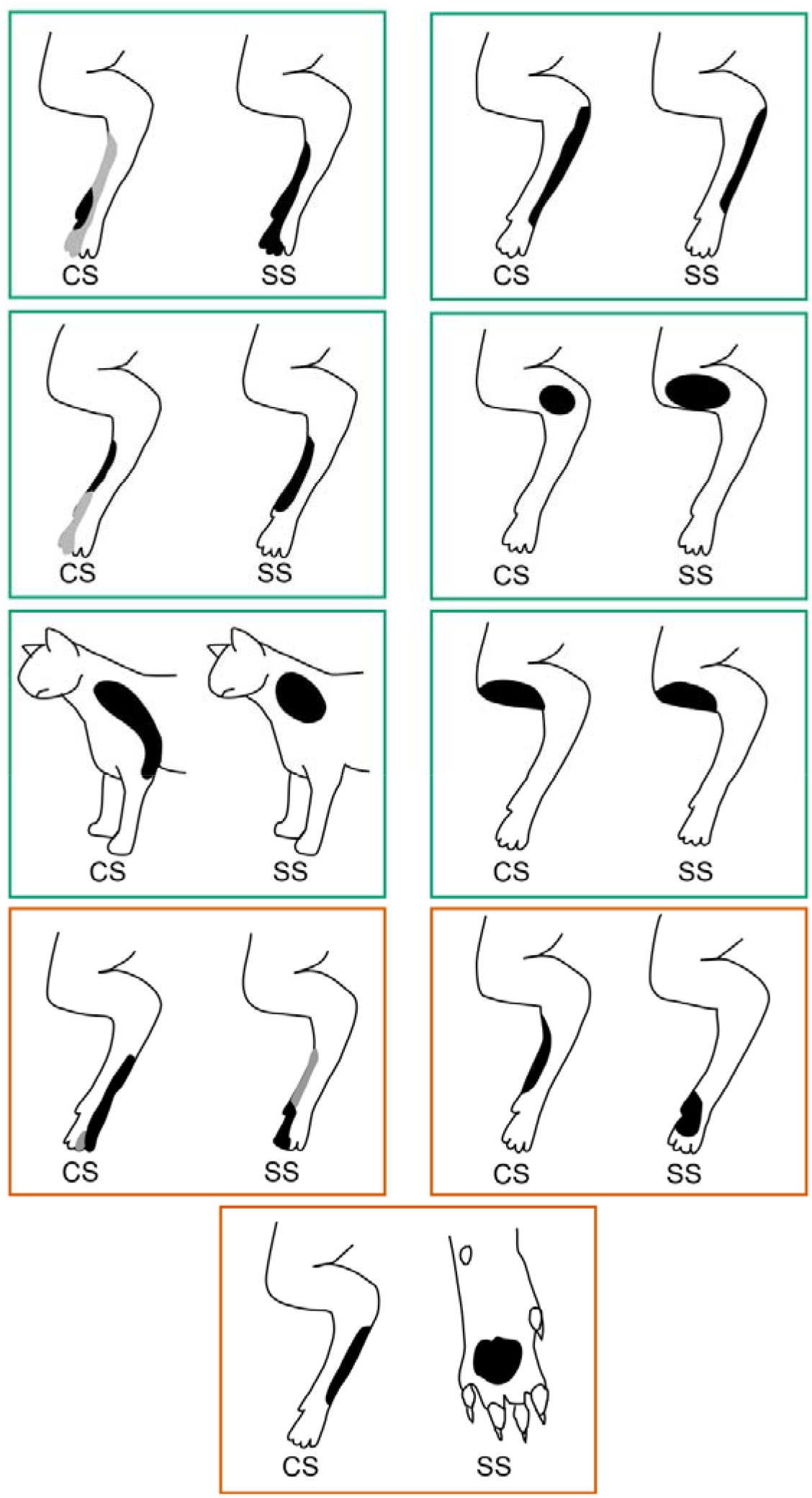
Receptive fields of complex spikes and simple spikes recorded from the same Purkinje cells. Complex spike and simple spike receptive fields for individual Purkinje cells (n=9) recorded when the animal was at rest. Pairs of receptive fields with the same receptive field classification are enclosed by green boxes; pairs with different receptive field classes are enclosed by red boxes. On each figurine black shading denotes regions of skin that generated the strongest responses, grey indicates total extent of receptive field. CS, complex spike; SS, simple spike.

### Pattern of activity of units with different receptive field classes during task performance

Of the 65 units grouped by class, 43 units (28 simple spikes, 6 complex spikes, 4 PCIs, and 9 mossy fibres) were recorded during task performance for a sufficient number of trials to permit quantitative analysis of neuronal activity during reaching, and also to examine the relationship between patterns of response and receptive field characterisation. The target for reach consisted of a Perspex tube containing a food reward and was located at approximately shoulder height of the animal (Fig. 1D). The cue to reach with the left forepaw was the opening of a door in front of the tube entrance.

The activity of almost all available units was clearly modulated at some point in the task, and the great majority exhibited strong modulations (a change in firing rate ≥ 2SD from baseline, see below). The largest peak of modulation was often accompanied by secondary, more subtle but distinct variations in discharge rate at other times in the task. This most likely reflected the fact that the task consisted of several key elements: (i) paw lift and forelimb reach, (ii) grasp, (iii) retrieval (including chewing and postural adjustments) and (iv) paw down. The complexity of the pattern of modulation was broadly related to location of peripheral receptive field. Units with distal (digit, paw or wrist receptive fields, classes 1-6 and 9) tended to show the most complex patterns of modulation i.e. tended to have 2 or 3 peaks of activity (Fig. 5A). They often changed their discharge rate before the time of paw lift or paw down, and also between the times of entry to and exit from the tube (when the fish morsel was grasped). By comparison, units with proximal or axial receptive fields (elbow, shoulder, neck, classes 8 and 10) tended to show simpler activity profiles (i.e. tended to have only one peak in activity, Fig. 5B).

**Figure 5.**
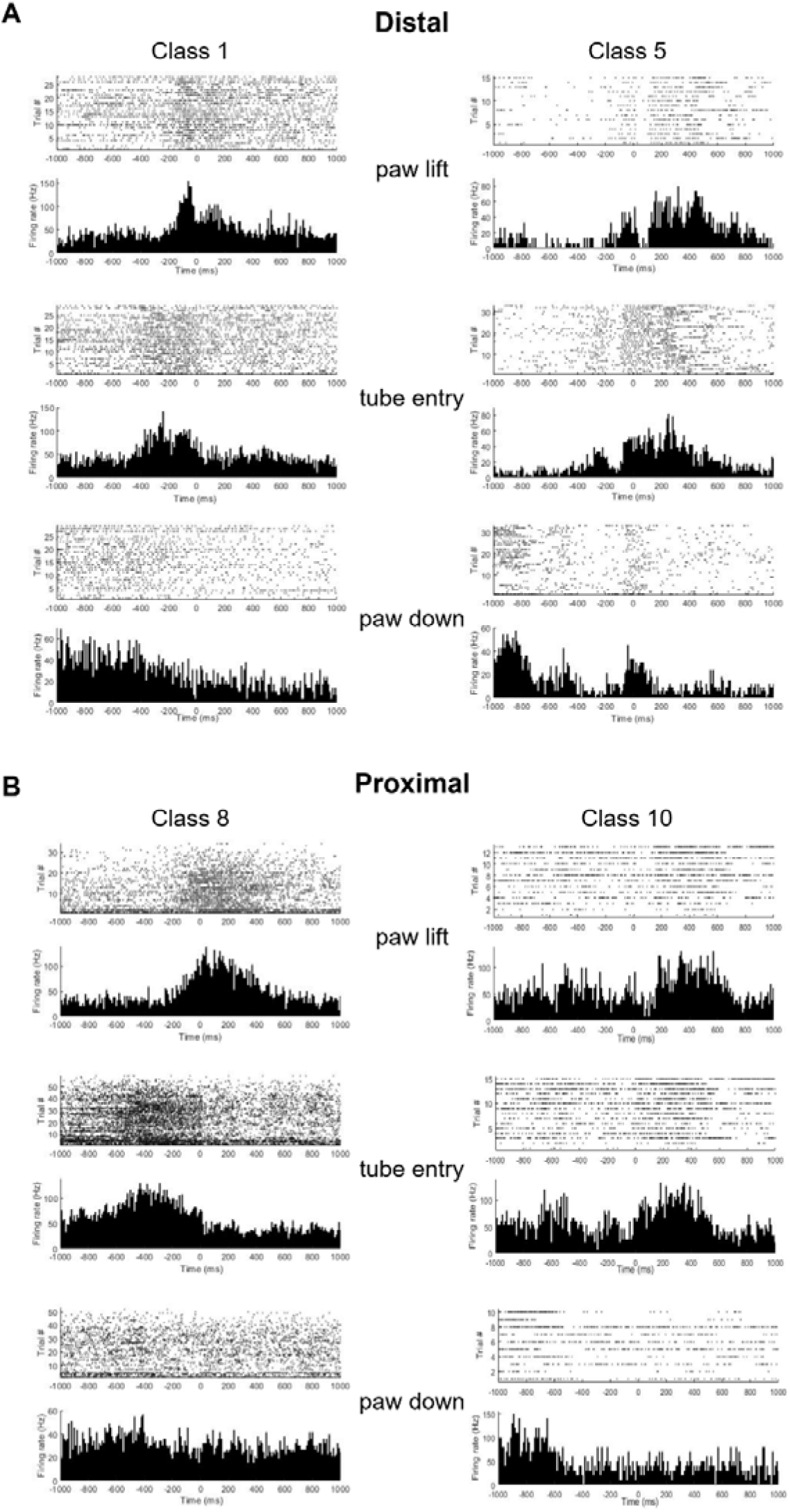
Pattern of activity during reaching of units with distal versus proximal limb receptive fields. **(A)** Example raster plots and peri-event time histograms (PETHs) for two classes of units (classes 1 and 5) with receptive fields located on the distal part of the ipsilateral forelimb. **(B)** Same as A but two example units (classes 8 and 10) with receptive fields located on the proximal part of the ipsilateral forelimb. In **A** and **B** top pair of panels shows rasters and PETHs with paw lift-off at time zero, middle pair of panels is tube entry at time zero and bottom pair of panels is paw touch-down at time zero.

Further analysis was confined to classes 1, 2, 6, 8, 9 and 10 (as each of these classes had ≥4 units), and also activity around the time of paw lift because this was a temporally precise task-related event and most units displayed modulation of their activity around this time. Figure 6A-F shows representative examples of the patterns of activity of units from classes 1, 2, 6, 8, 9 and 10 in relation to paw lift. Baseline firing rate between classes was not statistically significant (class 1 32.4 ± 15.5 Hz, n=8 units; class 2 22.6 ± 13.4 Hz, n=4 units; class 6 37.6 ± 7.2 Hz, n=5 units; class 8 31.6 ± 16.5 Hz, n=7; class 9 33.9 ± 30.1 Hz n=10 units; class 10 36.7 ± 21 Hz, n=4 units, Kruskal-Wallis p=0.934).

**Figure 6.**
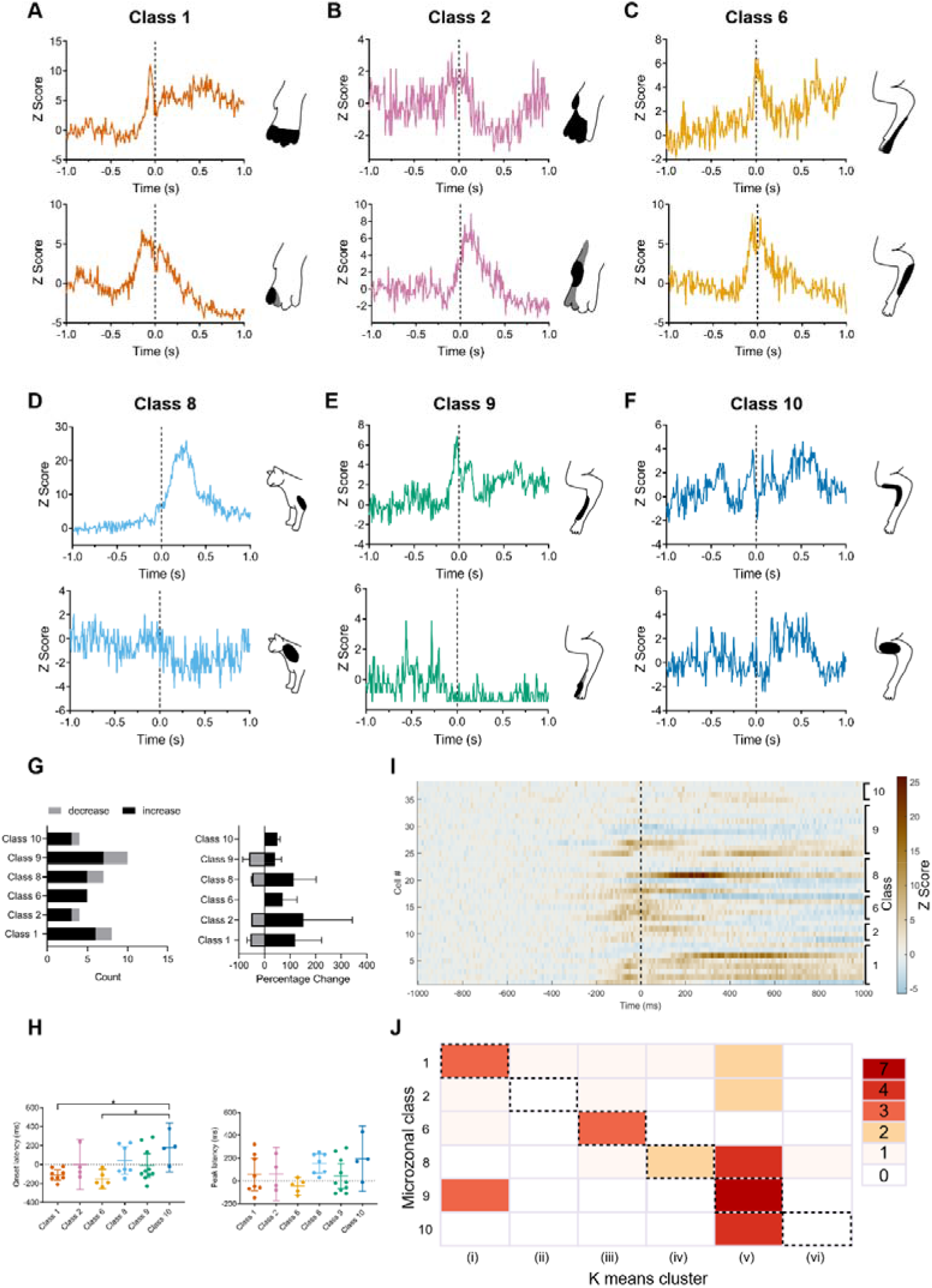
Individual patterns of activity during reaching of units within the same class. **(A-F)** Example normalized PETHs aligned to paw lift (time zero, vertical dashed line) for 12 units from 6 different receptive field classes. Inset shows receptive field location for each unit. Black shading denotes regions of skin that generated the strongest response; light grey areas denote total extent of receptive field. (**G**) Left panel, for each class the number of available units that displayed either an increase (black) or decrease (grey) in activity during paw lift-off. Right panel shows the average change in activity for the units that increased (black) or decreased (grey) their activity (mean± SD)**(H)** Left panel, onset latencies of significant change in unit activity per receptive field class in relation to paw lift-off (horizontal dotted line). Right panel, same as left panel but peak latency. Individual data points and mean± 95% confidence intervals, *p<0.05 **(I)** The activity patterns of units for which classes had ≥4 units (n=38, left hand y axis) shown as a heat map and arranged in relation to receptive class (right hand y axis). Each row indicates the activity pattern of a single unit aligned to paw lift-off (time zero, vertical dashed line). Firing rates are colour coded according to z-score values (colour scale to the right). **(J)** Confusion matrix showing the relationship between k-means clustering of the first three principal components of unit activity shown in (**I**) compared to grouping units based on receptive field class. The k-means clustering is divided into six columns (i-vi) plotted against six receptive field classes of individual units. The grid squares on the diagonal (highlighted with dashed borders) show the number of units where the k-means clustering matched receptive field class.

To quantify and compare changes in activity across different units, we first normalized firing rates as z-scores and defined changes in activity as statistically significant if they were ± 2 SDs from baseline (see Methods). Most units showed a significant increase in activity around the time of paw lift (Fig. 6G). Peak firing rates for units that increased or decreased compared to baseline were statistically significant (increase baseline=35.2 ± 24.5 Hz, peak=60 ± 42.1 Hz, n=30, p<0.0001; decrease baseline=23.8 ± 10.9 Hz, peak=11.7 ± 7.8 Hz, n=8, p=0.0078, Wilcoxon rank sum test, see also Supp. Fig. 2).

On average, the activity of all available units in class 1 and also class 6 exhibited significant changes in firing rate preceding paw lift (onset latency class 1, −110±63 ms, n=8, mean ± SD; class 6, −152 ± 83 ms, n=5; Fig. 6H). By comparison, the average activity of all units in class 2 and also class 9 showed a significant change in firing rate around the time of paw lift (onset latency class 2, 0 ± 165 ms, n=4; class 9, −8 ± 180ms, n=10, Fig. 6H). Class 8 and also class 10 units exhibited significant changes in firing after paw lift (onset latency class 8, 41 ± 159ms n=7; class 10, 180 ± 164 ms, n=4, Fig. 6H). The differences between onset latencies were statistically significant for classes 1 and 6 compared with class 10 (one-way ANOVA, p = 0.016, post hoc Tukey’s class 1 vs class 10 p=0.027, class 6 vs class 10 p=0.019, Fig. 6H). Peak latencies also differed between classes with those located distally occurring sooner than those classes located proximally, but this was not statistically significant (peak latency, class 1, 57.5 ± 171ms n=8; class 2, 60 ± 148ms n=4; class 6, −24 ± 64ms n=5; class 8, 151 ± 90ms n=7; class 9, 41 ± 152ms n=10; and class 10, 195 ± 181ms n=4, one-way ANOVA p=0.182; Fig. 6H).

To determine whether the pattern of neural activity between classes was related to differences in reaching performance, we used the duration of the reach (taken from time of paw lift to entry into the tube, see Methods) as a proxy of reaching kinematics. Reach duration was on average 320 ms ± 131 ms (n=7 cats, 2 animals only contributed to receptive field data). No statistical difference in reach duration was found between classes (Supp. Fig. 3, Kruskal Wallis p=0.56, n=6 classes). Similarly, no statistical difference was found between class, average firing rate and reach duration (ANCOVA p=0.585, n=38). Therefore, it seems reasonable to assume that the differences in pattern of neuronal activity observed between receptive field classes was not closely related to any variations in reach kinematics.

### Individual patterns of unit activity during reach

Inspection of the examples illustrated in Fig 6-A-F shows there was variation in individual unit response patterns belonging to the same receptive field class. The variation in activity in relation to paw lift across all available units is illustrated in the heat map shown in Figure 6I. In order to quantify the extent of consistency of the pattern of modulation of individual units within each class, we used principal component analysis (PCA) combined with k-means clustering (see Methods for details, see also Supp. Fig. 4). The number of clusters (n=6) was based on the number of receptive field classes available for analysis with a minimum of 4 units within each class (range 4-10 units per class).

For each unit, the comparison was visualized in a confusion matrix (Fig. 6J). The count in the diagonal (highlighted with dashed borders in Fig 6J) shows the number of units per class in which the k-means clustering matched receptive field class. Classes 6 and 9 had the greatest proportion of units that matched, whereas classes 2 and 10 had none, while classes 1 and 8 were intermediate in the proportion of matched units. Classes with a greater proportion of Purkinje cells had the best fit. Clustering was therefore also performed solely on the Purkinje cell population for classes which had ≥4 units (classes 1, 6, 8, and 9; see Supp. Fig. 5). To estimate the performance of the confusion matrix, we calculated the Adjusted Rand Index (ARI, see Methods for further details). The closer the ARI value to 100% the closer the fit, and a value over 95% is considered statistically significant. For the total population of available units used in Figure 6I, ARI = 96.6% while for the Purkinje cell population ARI= 98.6%. In sum, the response profiles of the individual units as determined by PCA were broadly related to receptive field class and this was most evident for Purkinje cells.

### Receptive field classes and cortico-nuclear projections

Within the C3 zone Purkinje cells belonging to the same subzonal cortical region have corticonuclear projections that converge on common territories within nucleus interpositus anterior (Apps and Garwicz, 2000). We therefore used receptive field classes as a frame of reference to sum the simple spike activity of different sets of Purkinje cells in order to estimate the pattern of modulation within the different cortico-nuclear pathways to nucleus interpositus anterior. As a first step, Figure 7A illustrates the population activity for all available Purkinje cells (n=26) recorded during task performance in relation to paw lift (hatched vertical line, Fig. 7A). The group PETH gives an estimate of the total pattern of simple spike modulation of Purkinje cells within the forelimb part of the C3 zone studied in the present experiments, and demonstrates that as a population, onset and peak of activity precedes paw lift, but activity remains elevated for ~600 ms afterwards (including paw entry to the Perspex tube at ~300 ms).

**Figure 7.**
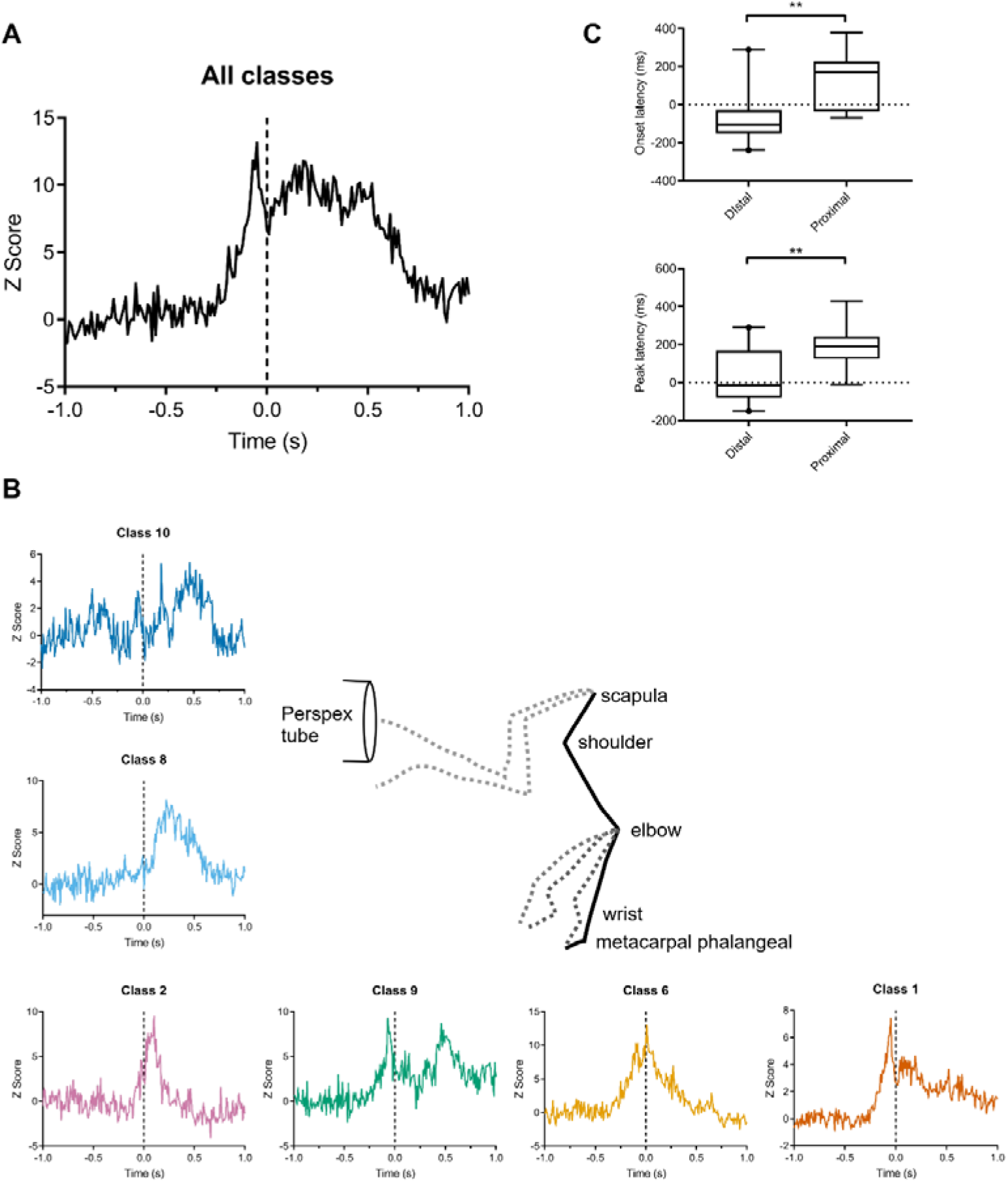
Summed pattern of activity during reaching for Purkinje cells within the same receptive field class. **(A)** Summed and z-scored PETH of Purkinje cell activity for all available receptive field classes aligned to paw lift (time zero, vertical dashed line, n=26 Purkinje cells). **(B)** Stick figure shows movement trajectory of different segments of the forelimb during reaching to a target. Summed and z-scored PETH of Purkinje cell activity for each available receptive field class aligned to paw lift (time zero, vertical dashed line). In clockwise direction the PETHs are in order of onset latency. **(C)** Onset and peak latencies of Purkinje cell activity for each class in relation to paw lift. Data are presented as box and whisker plots, central horizontal bar is the median, top and bottom horizontal lines of box indicate upper and lower quartiles. Whiskers indicate 95% confidence intervals. Values on whisker limits are shown as individual data points. ** p<0.01

The patterns of simple spike activity of Purkinje cells within each available receptive field class were also combined to produce a single PETH per class (Fig. 7B). Onset and peak latencies were grouped into proximal and distal location (Fig. 7C). There was a statistically significant difference in onset and peak latencies (Mann-Whitney onset latency U=147.5, p = 0.005, peak latency U=143.5, p=0.01; see Supp. Fig. 6 for onset latencies and peak activity relative to paw lift for each class).

The reaching component of the movement can be divided into two main phases: (i) paw lift, when there is initial flexion of the metacarpal phalangeal joints, followed by the elbow as the limb is raised from the surface (Martin et al., 1995). This initial phase coincides with the increases in population simple spike activity of Purkinje cells belonging to receptive field classes 1, 6 and 9 (lower row of PETHs in Fig. 7B). And (ii) the reach phase, involving (iia) dorsiflexion of the paw and then (iib) protraction of the shoulder and scapula as the limb is elevated towards the target. This second phase coincides with increases in population simple spike activity of Purkinje cells belonging to class 2 (iia) and classes 8 and 10 (iib, left hand column of PETHs in Fig. 7B). The results are therefore consistent with Purkinje cells belonging to different receptive field classes within the C3 zone collectively encoding different aspects of the reaching movement.

## Discussion

Our results demonstrate that the topographical organization of the C3 zone, originally established in anaesthetised and decerebrate preparations (Ekerot et al., 1991b; Garwicz et al., 1998; Ekerot and Jorntell, 2001), is also present in the awake animal. More specifically, our findings support the notion that the forelimb part of the C3 zone in the anterior lobe of the cerebellum has a detailed microzonal organization, which provides a structural framework for the neuronal information processing that underlies cerebellar control of skilled limb movements.

It should be emphasized that our mapping of receptive fields did not provide a comprehensive description at the microzonal level of resolution. Nonetheless, our results are compatible with such an organization. A similar limitation applies to the analysis of relationships between receptive fields and activity patterns of individual units, which was a central aim of the present study. Our results demonstrate that, during the performance of a reaching task, units belonging to different receptive field classes become active at different onset and peak latencies. By contrast, units belonging to the same class of receptive field displayed rather consistent activity patterns during behaviour, at least in some receptive field classes (e.g. classes 6 and 9). It remains to be determined whether the lower degree of consistency in activity patterns among units belonging to some receptive field classes (e.g. classes 8 and 10) was due to variability between units within the same microzone, or that, within the present sample, these classes contained a larger variety of individual microzones. Consistent with the latter possibility was the finding that units within classes 6 and 9 had less variation in receptive field location than those identified in classes 8 and 10.

### Simple spike and complex spike receptive fields of the same Purkinje cell

Previous in vivo studies have resulted in disagreement as to whether Purkinje cell simple spikes are activated by non-local groups of granule cells originating outside of a microzone (Ekerot and Jorntell, 2001) or from underlying, local granule cells (Bower and Woolston, 1983). Explanations for this difference include region of the cerebellum studied (paravermal C3 zone versus hemispheral crus I and II), species difference (cat versus rat), and experimental conditions. Our results are in keeping with more recent studies which have demonstrated that Purkinje cells can be activated by both local and distant granule cell inputs (Cramer et al., 2013; Valera et al., 2016). In our sample, two thirds of simple spikes and complex spikes recorded from the same Purkinje cells had very similar receptive field characteristics, implying local granule cell input, but the remainder differed markedly, implying non-local inputs. The simplest explanation that could account for the difference between our findings and those of Ekerot and Jorntell (2001) is that a decerebrate preparation was studied in the latter investigation.

Another possibility could be that the current study used animals that were well trained to perform the task. Training could modify the patterns of granule cell and/or parallel fibre inputs to cerebellar circuitry and result in plastic changes to microzonal maps. Valera et al., (2016) found that plasticity-inducing protocols could activate previously quiescent granule cell synapses, altering the pattern of connections between granule cells and Purkinje cells. Also, large changes in C3 zone receptive field maps have been reported by electrically stimulating parallel fibres in decerebrate cats (Jorntell and Ekerot, 2002). The stimulation induced changes to the receptive field size of Purkinje cell simple spikes, and in some cases, the skin overlying the complex spike receptive field which did not originally evoke a simple spike response, became the skin area that provided the strongest input to the simple spike receptive field. Future studies in awake animals will be needed to determine whether the receptive field characteristics of individual microzones can be altered by behavioural experience, for example during motor adaptation.

### Action-based organization and function

The patterns of spike activity generated during the reaching task by units within a receptive field class were in general qualitatively similar; and this was supported by correlation matrix analysis and the associated Adjusted Rand Index (Fig. 6). This was despite the fact that different types of units (e.g. Purkinje cell simple spikes, mossy fibres) were grouped together and that the analysis was carried out at the level of classes rather than microzonal subclasses. When the analysis was restricted to Purkinje cells within the same class, the Adjusted Rand Index was higher. This implies cerebellar cortical output is tuned in relation to receptive field identity.

In the present investigation, receptive fields of C3 microzones were similar to those found in previous studies in anaesthetised and decerebrate preparations in that their receptive field borders were close to joints and the areas from which the strongest activity was evoked were located eccentrically within the receptive field (cf. Ekerot et al., 1991b; Garwicz et al., 1998; Ekerot and Jorntell, 2001). Rather than a role in ‘sensory’ processing, the function of C3 microzones has been suggested to be related to the control of movements (Ekerot et al., 1991b). An increasing body of evidence suggests that the cerebellum is composed of a large number of olivo-corticonuclear ‘microcomplexes’, of which microzones and their associated receptive fields, constitute the cerebellar cortical component (Ito, 1984; Ekerot et al., 1991b; Garwicz et al., 1996; Garwicz et al., 1998; Apps and Hawkes, 2009). These microcomplexes are thought to represent the basic operational units of the cerebellum, with individual microcomplexes controlling ‘elemental’ movements, while collectively coordinating synergistic movements (Ekerot et al., 1995).

Multi-joint control of synergistic movements by the cerebellum is supported by many studies (Marple-Horvat and Stein, 1987; Fortier et al., 1989; Thach et al., 1992; Cooper et al., 2000; Becker and Person, 2019) and multi-joint movements have been shown to be particularly sensitive to cerebellar damage or experimental inactivation (Bastian et al., 1996; Topka et al., 1998; Cooper et al., 2000; Martin et al., 2000; Low et al., 2018; Becker and Person, 2019).

Microcomplexes have been suggested to coordinate movements either at the level of the cerebellar cortex through parallel fibres running across multiple microzones (Garwicz and Andersson, 1992; Apps and Garwicz, 2005), or by acting in concert on their target cerebellar nuclear cells (Ito, 1984). Previous studies have shown that groups of Purkinje cells firing synchronously can modulate their target cerebellar nuclear cells (Bengtsson et al., 2011; Blenkinsop and Lang, 2011; Person and Raman, 2011, 2012; Tang et al., 2019). However, anatomical tracing studies have yet to demonstrate the pattern of termination within the cerebellar nuclei of Purkinje cells located within an individual, physiologically-identified microzone.

To date, the highest resolution anatomical mapping approximates to studying the projections of different receptive field classes and has shown a detailed topography within cortico-nuclear projections (Garwicz and Ekerot, 1994; Garwicz et al., 1996; Apps and Garwicz, 2000; Ruigrok et al., 2008; Sugihara, 2011; Cerminara et al., 2013). Given the high degree of convergence in the corticonuclear projection (Apps and Garwicz, 2005), and the number of receptive fields in nucleus interpositus anterior is somewhat smaller than the number of microzones identified in the C3 zone (Garwicz and Ekerot, 1994) it is possible that microzones with similar receptive fields converge on common groups of nuclear neurons. Our study found that individual Purkinje cells could vary in their pattern of task-related activity, but when we pooled activity of Purkinje cells belonging to the same receptive field class (i.e. possessing similar receptive fields but potentially belonging to multiple microzones), we found systematic differences between classes in task-related activity. Purkinje cells with similar receptive fields may therefore act as an assembly on their target nuclear cells to control cerebellar output.

Microstimulation of nucleus interpositus has been shown to control multi-segmental muscle synergies (Ekerot et al., 1995). For example, stimulation of nuclear sites with receptive fields located on the ventral paw and forearm were associated with dorsiflexion of the wrist; stimulation of sites with receptive fields on the medial paw and forearm evoked elbow extension; stimulation of sites with receptive fields located on the lateral paw and forearm produced extension of the forearm; and stimulation of sites with receptive fields confined to the upper arm evoked shoulder flexion.

This functional activation can be mapped onto the results of the current study. For example, class 6 and 9 Purkinje cells, which have receptive fields located on the lateral and radial paw and forearm respectively had population activity that was related to forearm extension; class 2 with receptive fields on the paw had population activity that was related to dorsiflexion of the wrist; and class 8 and 10 with receptive fields located on the upper arm, had population activity related to shoulder flexion. Therefore, the microzonal organization of the C3 zone, previously suggested to be movement related (Ekerot et al, 1991b; Garwicz and Ekerot, 1994), and to provide error signals specific to different movement components (Garwicz 2002), appears to be matched by neuronal patterns of activity that reflect the control of different limb segments.

Taken together, the intrinsic organization of the cerebellar olivo-cortico-nuclear microcomplexes, their spino-olivo-cerebellar connections and the neuronal activity patterns during limb movements, are strongly suggestive of sensorimotor microcircuits operating in an action-based frame of reference (Apps and Garwicz, 2005). Action-based maps have also been found in the motor cortex, with different cortical sites responsible for ethologically relevant complex movements involving multiple muscles and joints (Graziano et al., 2002; Haiss and Schwarz, 2005; Graziano and Aflalo, 2007; Ejaz et al., 2015; Huber et al., 2020). A similar arrangement may apply to the cerebellum where populations of Purkinje cells within a subset of microzones with similar receptive fields combine to form a ‘super Purkinje cell’ to control its target group of cerebellar nuclear neurons (Apps et al., 2018) and co-ordinate multi-joint movements. We therefore believe it is essential when recording, analysing and interpreting activity patterns of single units in the paravermal part of the cerebellum, to take into consideration its microzonal organization.

The role of the cerebellum as a regulator, rather than an initiator or executor of movements, combined with the microcircuit organization and related activity patterns described here, suggest that the C3 zone controls forelimb movements by coordinating a relatively large number of muscle synergies (potentially corresponding to the number of microcomplexes), active at different times during skilled behaviour such as goal-directed reaching.

## Methods

All experimental and surgical procedures were performed in accordance with the UK Animals (Scientific Procedures) Act 1986 and were approved by the University of Bristol Animal Welfare and Ethical Review Body.

### Behavioural training

Nine purpose-bred adult male cats (4-6.5 kg) were trained non-aversively (ca. 6-8 weeks) to perform a reaching task. No aversive training techniques were used. The behavioural task was similar to that described previously (Alstermark et al., 1986; Apps et al., 1997). Briefly, the target for reach consisted of a Perspex tube (30 mm wide) placed in front of the animal at approximately shoulder height (Fig. 1D). The tube contained a small morsel of fish as a reward for successful completion of individual reaches. The cue to reach with the left forepaw was the opening of a door in front of the tube entrance. Throughout the sessions the animals were sitting in a natural seated posture and lightly restrained by a loosely applied harness. The head was unrestrained, which meant that the animal’s posture and movement were as natural as possible. At no point did any of the animals show any signs of discomfort. At the end of the recording session the animal was returned to a large communal pen where additional food was freely available.

### General surgical procedures

Following behavioural training, implantations were carried out using aseptic techniques and full surgical anaesthesia. Animals were first sedated with an intramuscular injection of medetomidine hydrochloride (Domitor, 15 μg/kg, Vetoquinol, UK) and the trachea intubated. Anaesthesia was induced with intra-venous propofol (Propofol Plus, 10mg/ml, Abbott Animal Health, USA) and maintained on gaseous isoflurane for the duration of the surgical procedure, except during periods of electrophysiological investigations during which anaesthesia was transferred back to propofol for cerebellar cortical mapping (see below). The concentration of isoflurane (1.5-3%) was adjusted during surgery in response to capillary O_2_ saturation, end tidal CO_2_, cardiac and respiratory rates and limb withdrawal reflex to pressure applied to the paws. Body temperature was monitored and maintained within physiological limits with the aid of a homeothermic heating blanket. At the end of surgery, a prophylactic injection of antibiotics (cefuroxime, Zinacef, 10mg/kg, GlaxoSmithKline, UK) was administered subcutaneously. Following surgery, the animals were monitored closely until they had recovered full consciousness. Analgesia was administered subcutaneously with buprenorphine (Buprecare, 20 mg/kg, Animal Care, UK) at the end of surgery followed by meloxicam (Metacam, 0.05 mg/kg, Boehringer Ingelheim, UK) for ~4-5 days post-surgery.

### Implantation procedure

Under general anaesthesia a craniotomy was made to expose the left anterior lobe paravermal cerebellum over lobule V, and a small recording chamber placed over the folia where the largest C3 zone-related electrophysiological responses were evoked by ipsilateral forelimb stimulation (see below). Up to 3 stainless-steel T-bolts were implanted to provide additional anchorage for the chamber and served as an indifferent for differential recording. For bipolar nerve stimulation, two pairs of Teflon-insulated wire were implanted around each of the left and right superficial radial nerves. A unipolar lead was implanted subcutaneously into the distal left forelimb to detect paw lift during reaching. Another lead was implanted in the back and served as an earth. All leads were fed subcutaneously from head-mounted micro-D connectors that were attached to the skull with dental acrylic cement, and a headpiece fashioned so as to incorporate the chamber.

### Cerebellar cortical mapping under general anaesthesia

Pairs of percutaneous stimulating needles were inserted temporarily into the ipsilateral and contralateral forelimbs and one or two brief (0.1ms, 1 kHz) square wave electrical pulses were delivered at an intensity to evoke a visible twitch in the stimulated limb. The responses to peripheral stimulation were recorded on the cerebellar surface using a lightly sprung ball electrode in order to identify the C zones of lobule V electrophysiologically. The resulting field potentials were amplified and bandpass filtered (0.03-5 kHz). Responses were classified according to previously defined response latencies and the presence of a contralateral response. Evoked fields in the C2 zone were defined as having a latency longer than 17ms in response to both ipsi- and contralateral stimulation whereas the neighbouring C1 and C3 zones were defined as having shorter (10-16ms) response latencies evoked by ipsilateral limb stimulation only (Ekerot and Larson, 1979; Trott and Armstrong, 1987; Apps and Garwicz, 2000).

### Recording arrangements and data acquisition in awake animals

Extracellular single-unit recordings confined to superficial cerebellar cortical layers were obtained in the awake animal during the reaching task using either parylene-insulated tungsten microelectrodes (WPI, impedance 2-5 MΩ) or a 16 channel Vector Array microelectrode (Neuronexus, Ann Arbour, Ml, USA). The microelectrodes were advanced using a stepping microdrive (NAN Instruments, Israel) mounted onto an x-y micromanipulator stage that was attached to the recording chamber during recording sessions. The x-y manipulator stage allowed the spatial co-ordinates of each electrode track to be precisely adjusted relative to the centre of the chamber. Signals were amplified and bandpass filtered (0.3-7.5 KHz). Paw lift-off from the foot plate and paw entry into the tube were continuously monitored via a high-frequency carrier signal applied to a copper contact plate and an IR beam located at the mouth of the tube, respectively (Fig. 1D). The opening of the door to the tube (the cue for the animal to reach) was monitored via a microswitch.

All neural and task event signals were captured using either customized Spike2 software running on a CED 1401Plus computer interface unit (Cambridge Electronic Design, Cambridge, UK) or a Cerebus Neural Signal Processor (Blackrock Microsystems LLC, USA).

### Experimental design

At the start of each recording session, the zonal location of the microelectrode was determined from the latency of the climbing fibre field potential evoked on electrical stimulation of each of the superficial radial nerves (Fig. 1B). Low intensity (non-noxious) paired-pulse stimulation was used, with the intensity increased until a small but visible limb twitch was observed. This was typically at an intensity twice the threshold (2T) for detecting a cerebellar cortical field response. Zonal identity was determined as described above.

Whilst the animal was sitting quietly, the microelectrode was advanced into the cerebellar cortex in small steps until a unit was isolated. Single units were identified by electrophysiological features of spontaneous and evoked neuronal activity and depth below the cerebellar surface. In order to ensure that we remained within the cortical zone identified at the start of the recording session, recordings were confined to the most superficial layers of the cerebellar cortex.

Purkinje cells were identified by the presence of complex spikes (Bloedel and Roberts, 1971; Armstrong and Rawson, 1979, Fig. 1C). In accordance with previous studies (Garwicz and Andersson, 1992) units that displayed a triphasic waveform and wide dynamic firing range, varying from silence to high frequency bursts at several hundred impulses per second were classified as mossy fibres. Units that could not be categorized as any of the above were classified as putative cortical interneurons.

Once a unit was isolated and identified, the receptive field for any peripheral somatic afferent inputs were determined and delineated by carefully delivering manual stimuli such as brushing of hairs, tapping of skin, palpation of muscle, and the passive movement of joints. No noxious stimuli were used. Receptive field classification was based on the detailed schema developed previously for the C3 zone in the anaesthetised cat (Ekerot et al., 1991a; Garwicz et al., 1998; Ekerot and Jorntell, 2001). The microzonal organization of the C3 zone was originally based on classification of climbing fibre receptive fields (Ekerot et al., 1991a), but was extended to the mossy fibre system (Garwicz et al., 1998) and comprises a total of 10 classes and approximately forty subclasses. The present study used this classification to group single units with similar receptive fields. Due to sampling constraints in the awake animal, receptive field grouping was confined to classes. The x-y coordinates of the micromanipulator stage allowed subsequent construction of topographical maps of receptive field identity as a function of spatial location within the C3 zone (Fig 2).

After receptive field mapping was complete, the animal began the task (described above) until they reached satiety. In some instances, the task was stopped midway and receptive field location checked to ensure the receptive field location remained stable.

### Spike sorting

Spikes were sorted with standard spike-sorting procedures off-line using either a clustering algorithm based on template matching and principal component analysis within Spike2 or with a semiautomated clustering algorithm (KlustaKwik; Kadir et al., 2014). The resulting clusters were manually inspected and refined with MClust (https://github.com/adredish/MClust-Spike-Sorting-Toolbox). Isolation of single units was verified by the presence of a distinct refractory period in the ISI histograms.

### Data analysis

Spike times and behavioural events (paw lift, tube entry) were analysed with custom-written scripts in MATLAB (v. 2018a). For each unit, perievent-time histograms (PETHs) of spike firing rates (bin width 10ms) were aligned to the onset of paw lift. To compare firing patterns across units, spike activity was transformed to z-scores using the baseline mean firing, where baseline was −1000 to – 500ms prior to the onset of movement. This baseline period avoided any preparatory changes in activity prior to movement. Changes in movement-related activity were considered significant when firing rates crossed ±2SD of the baseline mean. Onset latency was calculated to the leading edge of the first significant bin.

For Purkinje cells where both simple spike and complex spike receptive field maps could be reliably determined, the maps were superimposed in order to estimate the area of overlap. Area of overlap was expressed as the percentage of the total extent of the two receptive fields (each pair of black and grey shaded areas in Fig. 4 combined).

The duration of the reach was defined as the time interval between the paw lifting off the contact plate and entering the Perspex tube. Reach durations were used as a proxy to describe the reaching behaviour. Individual reaches were excluded from further analysis if they displayed reaction times of less than 150ms to the door opening, and/or if they displayed reach durations greater than 2.5 SD from the mean (Miller, 1991). The proportion of trials excluded based on these criteria was 12.5%.

We used a cluster-based method to determine whether units belonging to the same receptive field class had similar activity patterns. We first reduced the dimensionality of the standardised z-scored firing activity via principal component analysis. The coefficients associated with the first three principal components (which together explained >80% of the variance of the data, Forgy, 1965) were then used to calculate Euclidean distance. These data were then clustered using k-means with the number of clusters chosen to match the number of receptive field classes. This clustering grouped together units with similar activity patterns. A confusion matrix was used to describe the relationship between this k-means clustering of the data and grouping by receptive field class. To quantify how well the k-means clusters and receptive field class groups matched, we calculated the Adjusted Rand Index (Hubert & Arabie, 1985; Rand, 1971); since the k-means clustering reflected the neuronal pattern of activity this index measured the correspondence between activity pattern and receptive field class. To convert the Adjusted Rand Index into a statistic, the index was also calculated for simulated confusion matrices in which the receptive field class groups were assigned randomly; the values for 1,000 of these simulated confusion matrices were compared to the actual Adjusted Rand Index for the real confusion matrix. The closer the value of this statistic is to 100% the less likely any correspondence between receptive field class and activity patterns is by chance.

Data are expressed as mean ± SD unless stated otherwise. Statistical analyses were conducted using SPSS (IBM, USA). Normality was assessed using a Shapiro-Wilk test. Parametric and non-parametric tests were used accordingly.

### Histology

At the end of each experiment, animals were deeply anaesthetised with an intraperitoneal injection of pentobarbital. In some cases 200-400 nl of fluorescent-tagged material (20% solution of FluoroEmerald Molecular Probes, Eugene, OR, USA combined with green beads Lumafluor, New City, NY, USA) was injected into the cerebellar cortex (0.5–1 mm below the surface of the floor of the recording chamber) to mark the location where the majority of recordings where made (Fig. 1A). All animals were perfused transcardially with isotonic saline followed by 4% paraformaldehyde. The cerebellum was removed and stored in 4% paraformaldehyde at 4°C. The brain was transferred to a 30% sucrose solution after a week and allowed to sink. A freezing microtome was used to cut the cerebellum into 50 μm sagittal sections. Images were acquired using a Zeiss Axioskop II inverted microscope equipped with a CooLED pE-100 excitation system and Ocular digital monochrome camera and software.

## Supplementary Figures

**Figure supplement 1.**
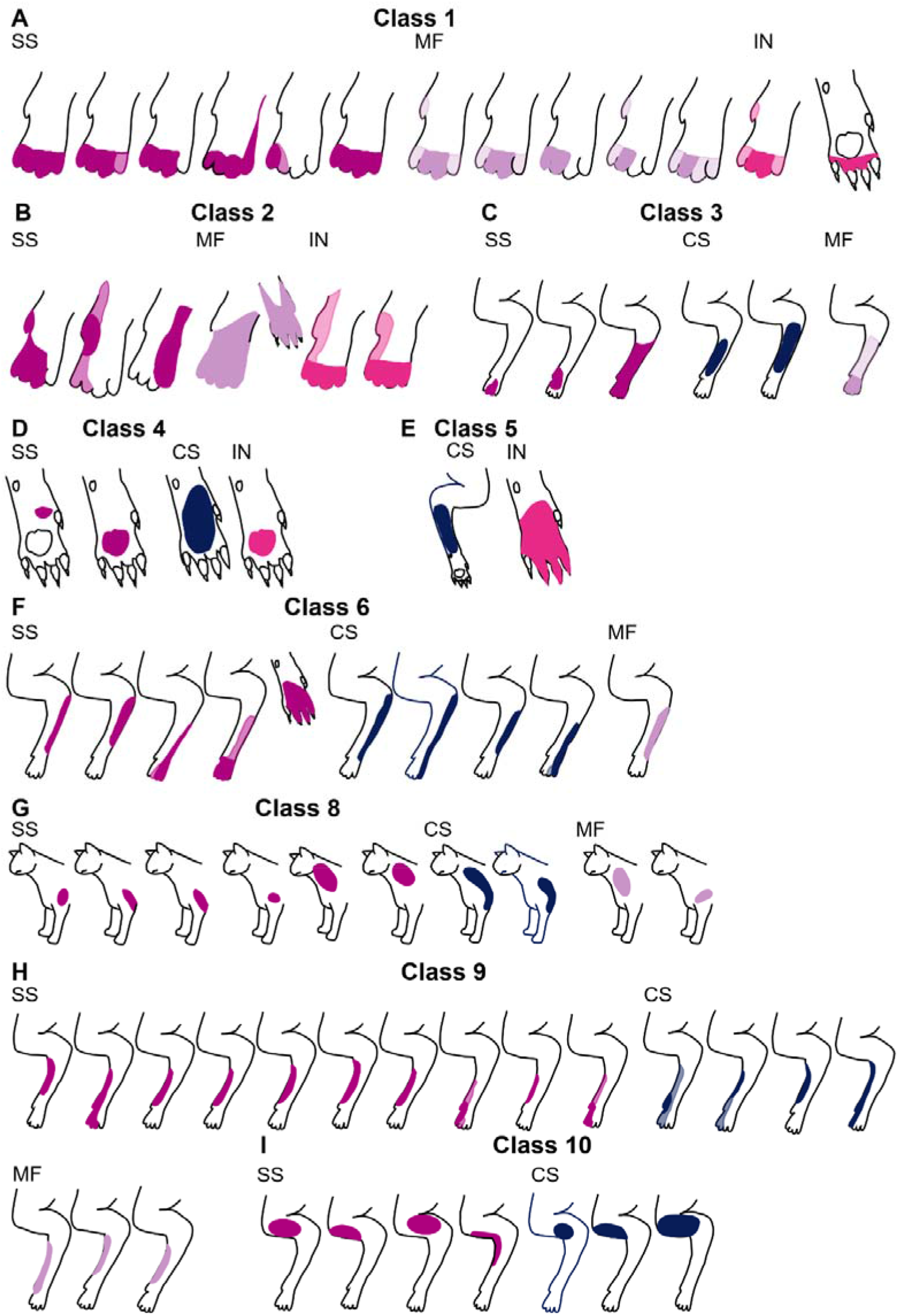
Classification of cerebellar units according to spatial organization of peripheral receptive fields. Units were categorised in accordance with previous classification of Purkinje cell complex spike (Ekerot et al., 1991a) and mossy fibre receptive field characteristics (Garwicz et al., 1998). Each figurine shows the receptive field of a single unit organized in relation to receptive field class and subdivided into those units identified as Purkinje cell simple spikes (SS, purple), complex spikes (CS, dark blue), mossy fibres (MF, light purple) or putative cortical interneurons (IN, pink). (**A**) Class 1 (n=13); (**B**) Class 2 (n=6); (**C**) Class 3 (n=6); (**D**) Class 4 (n=4); (**E**) Class 5 (n=2); (**F**) Class 6 (n=9); (**G**) Class 8 (n=10); (**H**) Class 9 (n=17); (I) Class 10 (n=7). No units were found for class 7. On each figurine darker shading denotes regions of skin that generated the strongest response; lighter shading denotes total extent of receptive field.

**Figure supplement 2.**
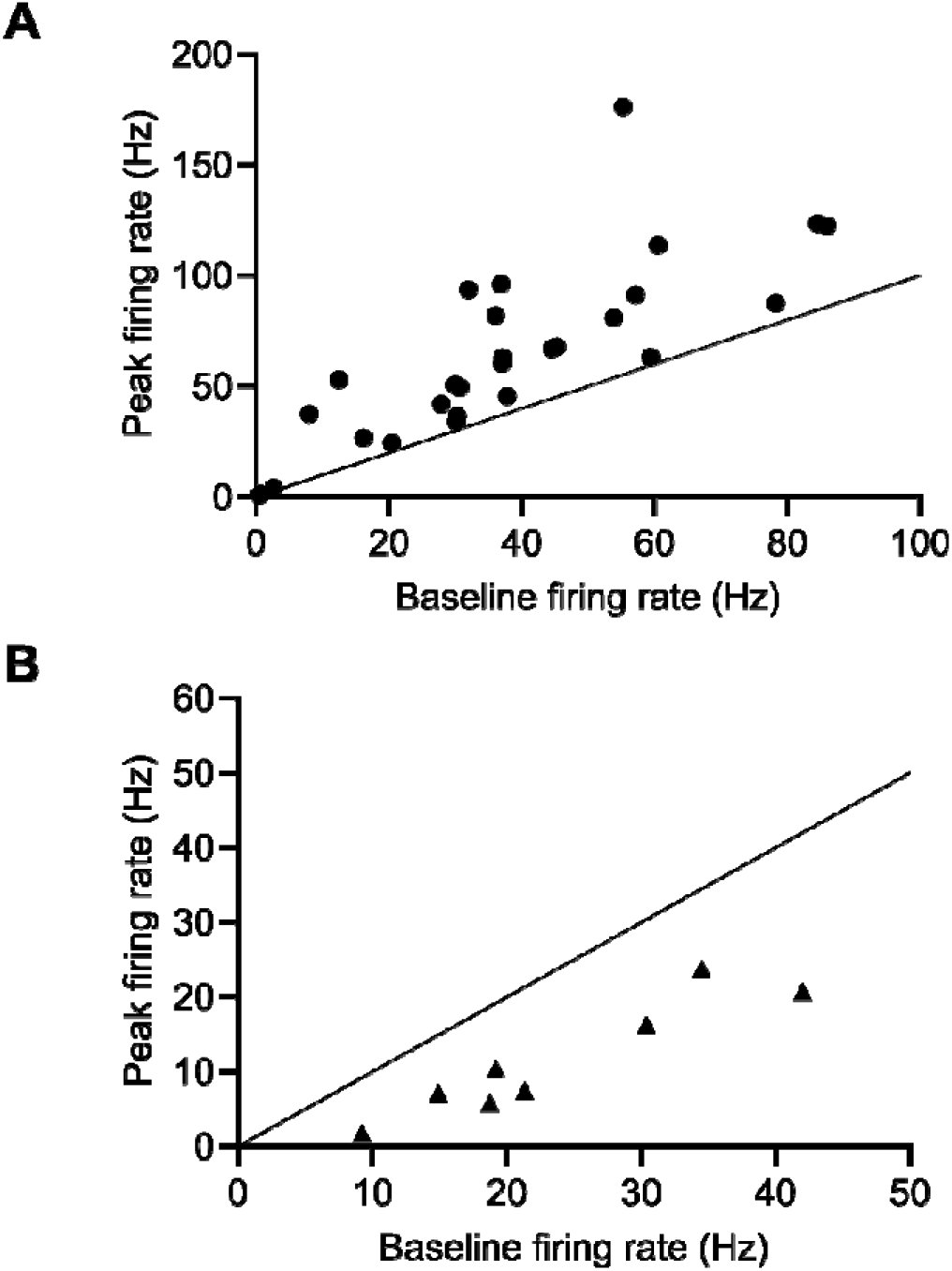
Changes in firing rate during paw lift. Baseline vs peak firing rates for all units which either increased (**A**) or decreased (**B**) their activity during reach. Diagonal black line of equivalence.

**Figure supplement 3.**
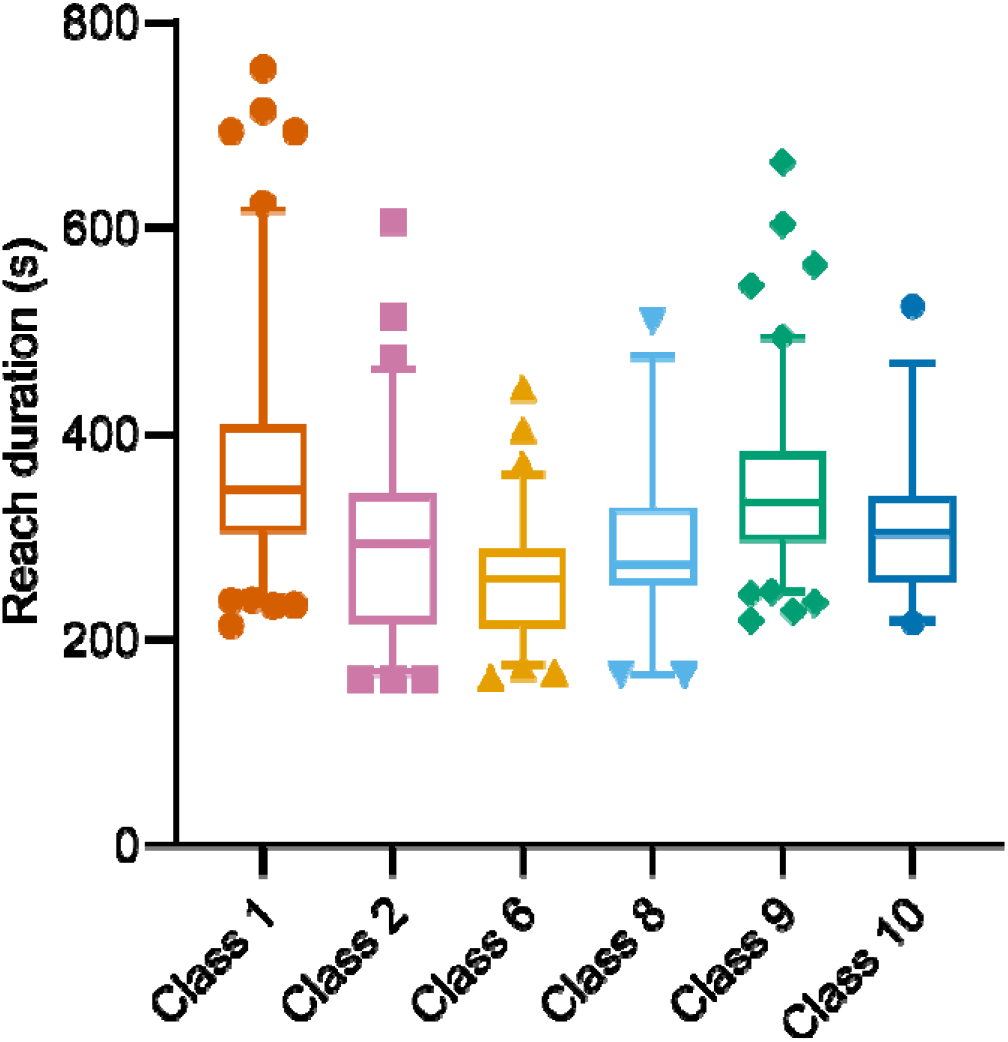
Box and whisker plots showing reach duration for units from each receptive field class. In each plot, the central horizontal bar is the median, top and bottom horizontal lines of box indicate upper and lower quartiles. Whiskers indicate 95% confidence intervals. Points above and below the whiskers are shown as individual data points.

**Figure supplement 4.**
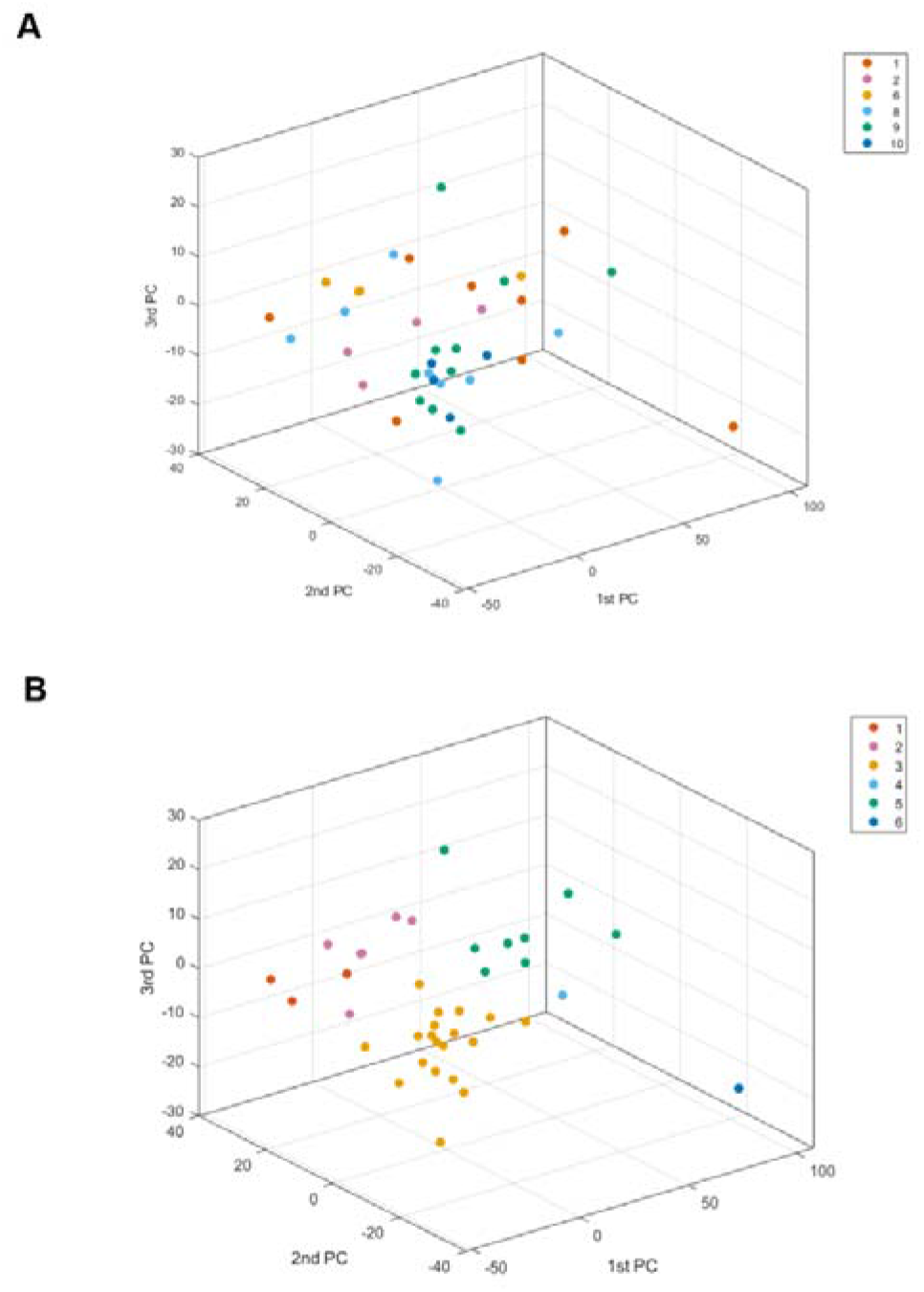
Clustering of units. (**A**) Projection in 3 dimensions of the first three principal components of the normalized z-scored spike firing rates for individual units. Data points colour coded according to receptive field class. (**B**) k-means clustering of the first three principal components. The number of clusters (n=6) was based on the number of receptive field classes with at least 4 available units to run the comparison. Data points colour coded according to k-means cluster.

**Figure supplement 5.**
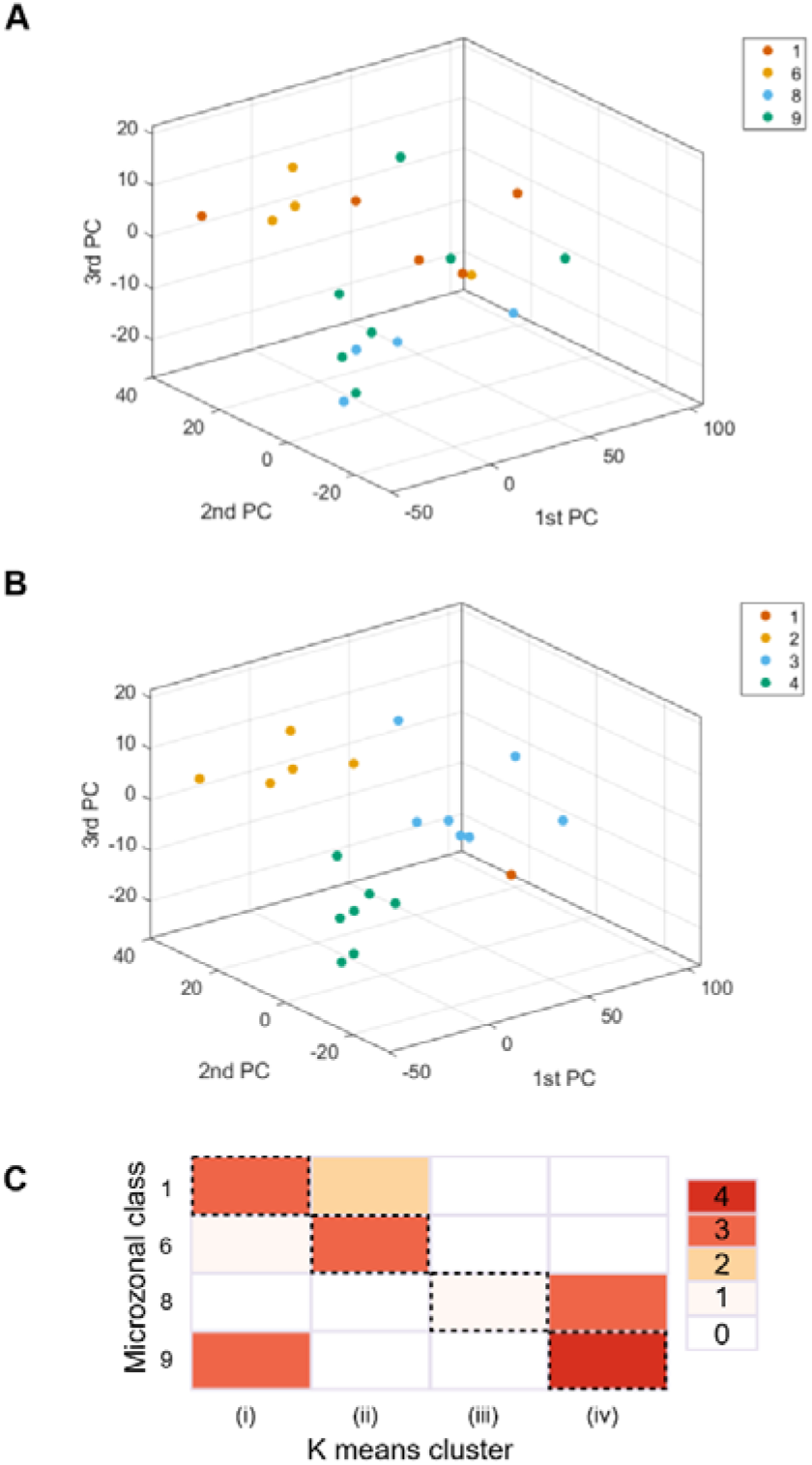
Clustering of Purkinje cells. (**A**) Projection in 3 dimensions of the first three principal components of the normalized z-scored spike firing rates for individual Purkinje cells categorised by receptive field to belong to one of four classes (colour coded 1, 6, 8 or 9). (**B**) K-means clustering of the first three principal components. The total number of clusters (colour coded 1-4) was based on the number of available receptive field classes. **(C)** Confusion matrix showing the relationship between k-means clustering of the first three principal components of unit activity shown in (**B**) compared to grouping units based on receptive field class. The k-means clustering is divided into four columns (i-iv) plotted against four classes of individual units. The grid squares on the diagonal show the number of units where the k-means clustering matched receptive field class.

**Figure supplement 6.**
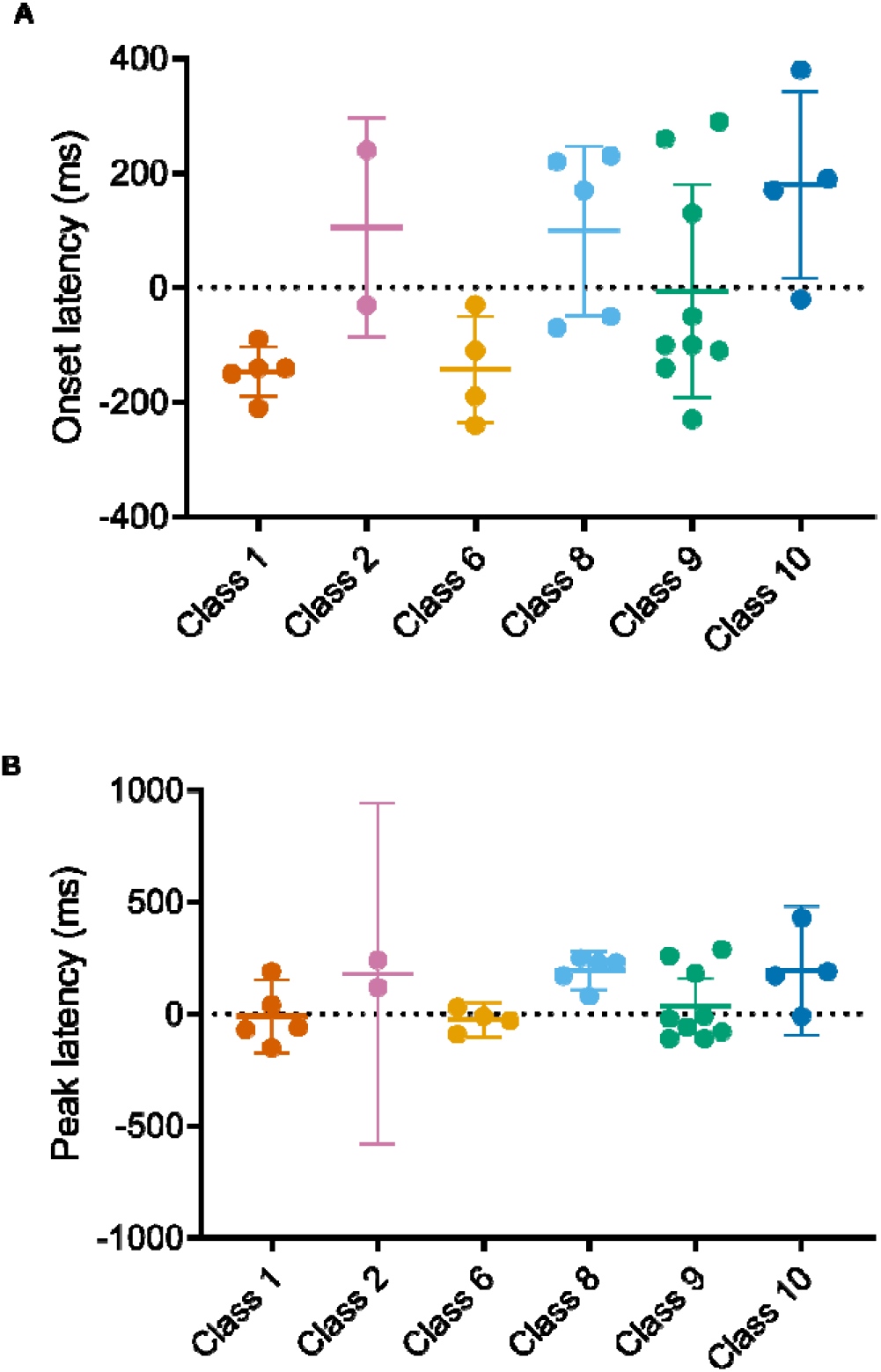
Latencies of Purkinje cell activity in relation to receptive field class. **(A)** Onset latency of significant change in unit activity per receptive field class in relation to paw lift. (**B**). Same as A but peak latency. Data are presented as mean± 95% confidence intervals.

## Acknowledgements

This work was supported by the Medical Research Council UK (G1100626) to NLC and RA. HD was supported by a BBSRC SWBio DTP Award (1503834). The authors are grateful for the support and assistance of Dr Jo Murrell (veterinary anaesthetist) and Rachel Bissett.

